# Nitroxide functionalized antibiotics are promising eradication agents against *Staphylococcus aureus* biofilms

**DOI:** 10.1101/579896

**Authors:** Anthony D. Verderosa, Rabeb Dhouib, Kathryn E. Fairfull-Smith, Makrina Totsika

## Abstract

Treatment of *Staphylococcus aureus* biofilm-related infections represents an important medical challenge worldwide, as biofilms, even of drug-susceptible *S. aureus* strains, are highly refectory to conventional antibiotic therapy. Nitroxides were recently shown to induce dispersal of Gram-negative biofilms *in vitro*, but their action against Gram-positive bacterial biofilms remains unknown. Here we demonstrate that the biofilm dispersal activity of nitroxides extends to *S. aureus*, a clinically important Gram-positive pathogen. Co-administration of the nitroxide CTEMPO with ciprofloxacin significantly improved the antibiotic’s biofilm-eradication activity against *S. aureus*. Moreover, covalently linking the nitroxide to the antibiotic moiety further reduced ciprofloxacin’s minimal biofilm eradication concentration. Microscopy analysis revealed that fluorescent nitroxide-antibiotic hybrids could penetrate *S. aureus* biofilms and enter into cells localising at the surface and base of the biofilm structure. No toxicity was observed for the nitroxide CTEMPO and the nitroxide-antibiotic hybrids against human cells. Taken together, our results show that nitroxides can mediate dispersal of Gram-positive biofilms and that dual-acting biofilm-eradication antibiotics could provide broad-spectrum therapies for the treatment of biofilm-related infections.

## INTRODUCTION

*Staphylococcus aureus* is a Gram-positive commensal and opportunistic human pathogen, which is a major cause of nosocomial and community-acquired infections (1). The inherent ability of *S. aureus* to attach to medical devices and host tissues and establish biofilms is a major driver of failing antibiotic therapies and the persistence of chronic infections (2-4). Thus, there is an urgent need for novel strategies for the treatment and eradication of *S. aureus* biofilms.

Current antimicrobial strategies, which are effective against planktonic bacteria, often have little or no effect when administered to biofilms (5, 6). During infection, bacteria reside mostly within biofilms but can revert to the planktonic lifestyle by modulating the expression of specific genes (7). Consequently, the development of small molecules with the ability to trigger a cell change from biofilm to the planktonic state has become a promising area of anti-biofilm research (8, 9).

One of the most promising small molecules with anti-biofilm activity is nitric oxide (NO), which has been shown to inhibit biofilm formation and trigger biofilm dispersal (10-12) in a variety of biofilm-forming bacteria (13). In *Pseudomonas aeruginosa*, for example, treating a mature biofilm with sub-lethal NO concentrations (pM to nM range) triggers a transition from the sessile (biofilm) to the motile (planktonic) state, providing a means of effective antibiotic therapy (11, 13). The anti-biofilm properties of NO in *P. aeruginosa* are mediated by regulation of intracellular levels of the secondary messenger cyclic dimeric guanosine monophosphate (cyclic di-GMP), which plays a pivotal role in biofilm development; high levels facilitate biofilm formation, while low levels prompt biofilm dispersal (7, 14). However, the targeted and controlled delivery of NO to biological systems as therapies is a challenge due to the reactive nature and short half-life (0.1 – 5 seconds) (15) of the gaseous molecule. Consequently, methods for circumventing this challenge have been the focus of intensive research over the past several years, with the use of NO-donors in biofilm dispersal now comprehensively documented (16, 17). A variety of NO-donor compounds with anti-biofilm activity have already been described. However, donor molecules are often themselves inherently unstable (18), necessitating an alternative approach.

Nitroxides are long-lived, stable free radical species, which contain a disubstituted nitrogen atom linked to a univalent oxygen atom (19). Nitroxides are considered sterically hindered, structural mimics of NO, as both compounds contain an unpaired electron, which is delocalized over the nitrogen-oxygen bond. The biological activity of nitroxides is often attributed to their NO-mimetic properties, with both being efficient scavengers of protein-derived radicals (20). Nitroxides, however, are mostly air-stable crystalline solids, in contrast to NO, which is unstable and gaseous at room temperature. This significant difference makes nitroxides ideal candidates for circumventing the handling and delivery issues associated with NO.

We have previously demonstrated the ability of nitroxides to inhibit and disperse bacterial biofilms of *P. aeruginosa* and *E. coli* (21, 22). Nitroxides were also shown to enhance biofilm antibiotic susceptibility when co-administered as a combined treatment (23). Furthermore, we have shown that by covalently tethering a nitroxide to the antibiotic ciprofloxacin and thus delivering the antibiotic at the site of nitroxide mediated dispersal, we could achieve effective eradication of mature *P. aeruginosa* biofilms (24, 25). While the dispersal and anti-biofilm properties of nitroxides have been successfully documented against Gram-negative bacteria, their anti-biofilm properties against Gram-positive bacteria remain unexplored. Here we have employed a reproducible, high-throughput *in vitro* biofilm assay to comprehensively examine the full antimicrobial, anti-biofilm, and biofilm-eradication potential of nitroxides and antibiotic-nitroxide hybrids against *S. aureus*. Our findings demonstrate the anti-biofilm action of nitroxides against a clinically important Gram-positive pathogen extending this promising therapeutic strategy to a large number of *S. aureus* biofilm-related infections.

## RESULTS

### The nitroxide CTEMPO can disperse established *S*. *aureus* biofilms

To examine the biofilm dispersal properties of the nitroxide 4-carboxy-2,2,6,6-tetramethylpiperidin-1-yloxyl (CTEMPO) against *S. aureus* in a high-throughput system, we modified the static Calgary Biofilm Device (CBD) in a way that allows the specific eradication of disperser cells with antibiotics (ciprofloxacin). This methodology ensured that *S. aureus* cells were immediately eliminated after dispersing from the biofilm (ciprofloxacin susceptible) preventing them from reverting to biofilm growth (ciprofloxacin resistant). This modification allowed us to monitor biofilm dispersal in a static biofilm system similar to that achieved in a dynamic flow cell system, where media influx physically removes disperser cells. In our system, this was achieved by the addition of 6 µM ciprofloxacin (minimum bactericidal concentration (MBC) against disperser *S. aureus* ATCC 29213 cells) in every well treated with the nitroxide CTEMPO at a concentration range.

Using this methodology, established *S. aureus* ATCC 29213 biofilms were co-treated with CTEMPO (160 – 2.5 µM) and ciprofloxacin (6 µM) or ciprofloxacin alone (6 µM) for 24 hours. Viable bacteria remaining in the treated biofilms were recovered in media without nitroxide or antibiotics and monitored for growth, with lag time recorded as a direct measure of the initial colony forming units (CFU) present in each well. Significant increase in lag time, indicative of a significant reduction in biofilm-associated bacteria recovered post-treatment, was observed at CTEMPO concentrations in the range of 10 to 80 µM (*p*= 0.0003, Kruskal-Wallis test), compared to control biofilms treated with ciprofloxacin alone (Figure 1). As no antimicrobial activity was observed for CTEMPO alone at those concentrations against planktonic or biofilm-residing *S. aureus* ATCC 29213 cells (MIC >3200 µM, MBEC >1200 µM; Table 1), the nitroxide was most likely inducing cell dispersal in biofilms, not bacterial killing. CTEMPO alone (no ciprofloxacin) had no impact on the number of viable bacteria recovered from the biofilm.

**Figure 1:**
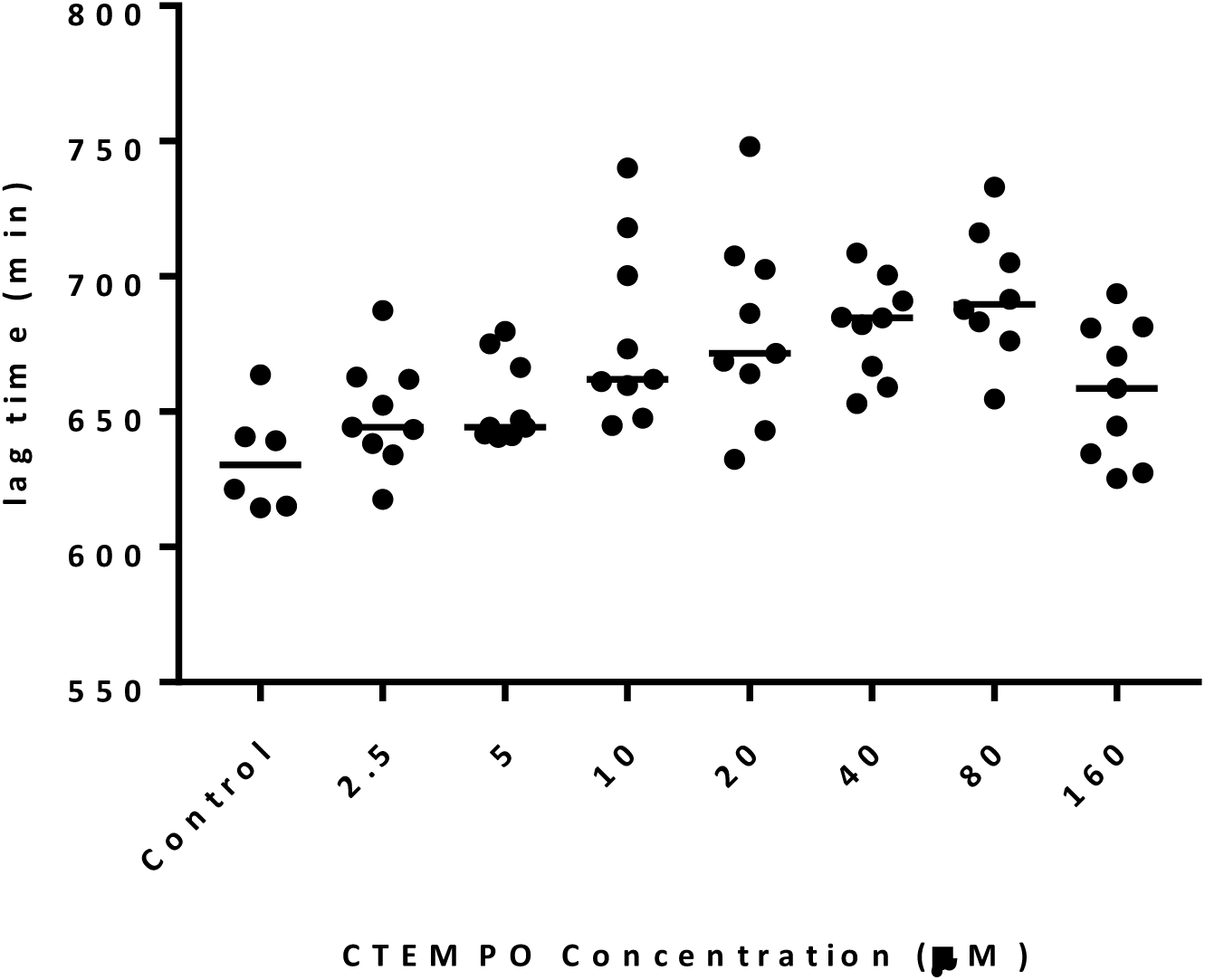
Nitroxide-mediated *S. aureus* ATCC 29213 biofilm dispersal. Established *S. aureus* biofilms were treated with the nitroxide CTEMPO (at a concentration range of 160 – 2.5 µM) and ciprofloxacin (6 µM) or with ciprofloxacin alone (6 µM) (Control) for 24 hours. Biofilm-associated bacteria were recovered after treatment and enumerated as described in methods. Lag time (min) from growth curves of recovered bacteria was calculated using nonlinear regression. Dot plots show data from treated and control biofilms obtained from 3 biological repeats performed with at least 3 technical replicates. Lines show group medians. Group medians were compared using the Kruskal-Wallis test (*p*= 0.0003).

**Table 1:** MIC and MBEC values for ciprofloxacin, CTEMPO, co-treatment (ciprofloxacin/CTEMPO), and antibiotic-nitroxide hybrids against *S. aureus* ATCC 29213.

|  | Treatment | MIC<sup>1</sup> (μM) | MBEC<sup>2</sup> (μM) |
| --- | --- | --- | --- |
| **Controls** | Ciprofloxacin | ≤1.5 | ≤4096 |
|  | CTEMPO | >3200<sup>[a]</sup> | >1200 |
| **Co-treatment of *S. aureus* biofilms** | Ciprofloxacin & CTEMPO (2.5 μM) | nd | ≤4096 |
|  | Ciprofloxacin & CTEMPO (8 μM) | nd | ≤256 |
|  | Ciprofloxacin & CTEMPO (10 μM) | nd | ≤512 |
|  | Ciprofloxacin & CTEMPO (20 μM) | nd | ≤512 |
|  | Ciprofloxacin & CTEMPO (40 μM) | nd | ≤512 |
|  | Ciprofloxacin & CTEMPO (80 μM) | nd | ≤2048 |
| **Hybrid treatment of *S. aureus* biofilms** | Cipro-PROXYL | ≤6.3 | ≤1024 |
|  | Cipro-TEMPO | ≤12.5 | >1024<sup>[a]</sup> |
|  | Cipro-TMIO | ≤1.5 | ≤64 |
|  | Fluoro-TEMPO | ≤20 | ≤720 |
|  | Fluoro-TEMPOMe | >1200<sup>[a]</sup> | >1480<sup>[a]</sup> |

<sup>1</sup>MICs were determined via the broth microdilution method in accordance with CLSI
standard. <sup>2</sup>MBECs were determined using the CBD. [a] Highest concentration tested. nd: Not
done.

### CTEMPO co-administration improves the efficacy of ciprofloxacin against *S*. *aureus* biofilms

As nitroxides appear to induce *S. aureus* biofilm dispersal, we sought to examine whether this could improve the efficacy of ciprofloxacin against established *S. aureus* biofilms. Ciprofloxacin is a commonly prescribed fluoroquinolone antibiotic with potent antimicrobial activity against planktonic *S. aureus* ATCC 29213 (26). We confirmed the MIC of ciprofloxacin for this strain to be within the previously published range (1.5 - 0.4 µM; Table 1) (26). Despite the low MIC, established *S. aureus* ATCC 29213 biofilms required 4096 µM of ciprofloxacin for complete (99.9%) eradication (concentration range tested: 4096 - 32 µM), demonstrating the biofilm’s extremely high antibiotic tolerance. The minimum biofilm eradication concentration (MBEC) of ciprofloxacin against established *S. aureus* ATCC 29213 biofilms was thus determined to be ≤4096 µg/mL (Table 1).

Co-administration of CTEMPO (at 2.5, 8, 10, 20, 40, or 80 µM) with ciprofloxacin (at a concentration range of 4096 - 32 µM) against established *S. aureus* ATCC 29213 biofilms resulted in a significant reduction in the MBEC value of ciprofloxacin at CTEMPO concentrations between 8 - 80 µM (Table 1). Ciprofloxacin potentiation was greatest with 8 µM CTEMPO co-administration, which reduced ciprofloxacin’s MBEC value from ≤4096 µM to ≤256 µM, representing a 16-fold improvement in the drug’s efficacy against Gram-positive *S. aureus* ATCC 29213 biofilms (Table 1). Taken together, these results demonstrate the ability of nitroxides to significantly improve the efficacy of antibiotic therapy against *S. aureus* biofilms.

### Ciprofloxacin-nitroxide hybrids are potent *S*. *aureus* biofilm-eradication agents

Considering the synergistic effect of nitroxide-ciprofloxacin co-administration on *S. aureus* biofilm eradication, we evaluated whether combining the nitroxide and antibiotic within one molecule would offer greater improvement. We rationalized that by localising the antibiotic directly at the site of nitroxide-mediated dispersal, it could more effectively eradicate dispersed cells than in combination treatment. To test this hypothesis, we utilized several ciprofloxacin-nitroxide hybrids that we have previously generated (Figure 2) (25, 27). Cipro-PROXYL, cipro-TEMPO, and cipro-TMIO (Figure 2) were first screened in MIC assays for activity against planktonic *S. aureus* ATCC 29213 (Table 1). All compounds were determined to possess potent *S. aureus* ATCC 29213 activity (MIC range 1.5 - 12.5 µM) with cipro-TMIO being the most active (MIC ≤1.5 µM, same MIC as ciprofloxacin). Subsequent minimum bactericidal concentration (MBC) analysis of cipro-TMIO and ciprofloxacin revealed that cipro-TMIO was at least twice as bactericidal as ciprofloxacin against *S. aureus* ATCC 29213 cells (cipro-TMIO MBC: ≤1.5 µM vs. ciprofloxacin MBC: ≤3.0). Then, established *S. aureus* ATCC 29213 biofilms were treated with cipro-PROXYL, cipro-TEMPO, cipro-TMIO, fluoro-TEMPO, or fluoro-TEMPOMe (fluoro-TEMPO derivative with free radical removed). Cipro-PROXYL, cipro-TMIO, and fluoro-TEMPO all exhibited potent biofilm-eradication activity (Table 1). However, cipro-TMIO was by far the most potent agent with an MBEC value of ≤64 µM. This made cipro-TMIO at least 4-fold more potent than ciprofloxacin/CTEMPO co-treatment (MBEC ≤256 µM) and >64-fold more potent than ciprofloxacin alone (MBEC ≤4096 µM) against *S. aureus* biofilms. These results demonstrate the advantage of covalently linking a nitroxide moiety to an antibiotic to enhance its activity against Gram-positive biofilms.

**Figure 2:**
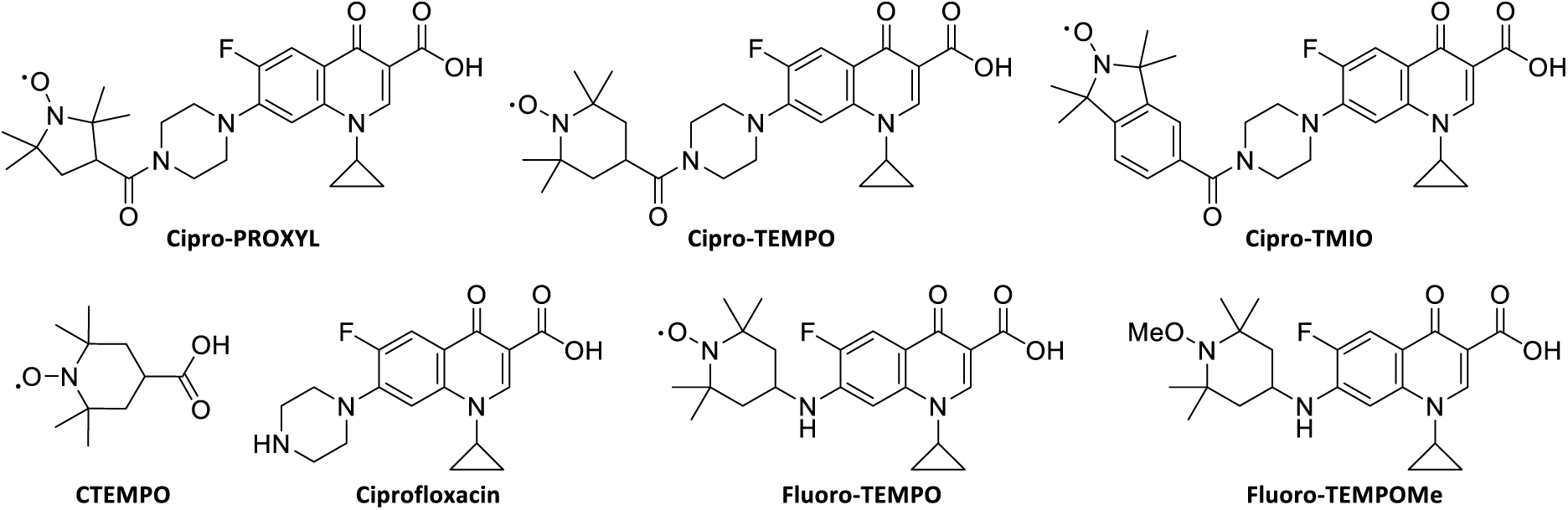
Chemical structures of CTEMPO, ciprofloxacin, cipro-PROXYL,(25) cipro-TEMPO,(25) cipro-TEMPO,(25) fluoro-TEMPO,(27) and fluoro-TEMPOMe (27).

### Fluoroquinolone-nitroxide hybrids can penetrate *S*. *aureus* biofilms and enter into surface and base-residing cells

To investigate the impact of nitroxide-antibiotic hybrids on established *S. aureus* biofilms in more detail, we utilized the recently reported profluorescent fluoroquinolone-nitroxide hybrid fluoro-TEMPO (27). Fluoro-TEMPO was found to be most active against *S. aureus* and emit bright fluorescence upon cell entry (27), making it a valuable switch-on fluorescent probe for studying *S. aureus* biofilm eradication by nitroxide-antibiotic hybrids. We first established the MBEC value for fluoro-TEMPO (MBEC ≤720 µM; Table 2) and found it to be at least 5-times more potent than ciprofloxacin treatment of *S. aureus* ATCC 29213 biofilms, further supporting the increased potency of tethered nitroxide-antibiotic hybrids. Importantly we showed that this effect was specific to the presence of the free radical nitroxide as the methoxyamine derivative fluoro-TEMPOMe had lost its antibacterial activity (MIC >1200 µM and MBEC >1480 µM; Table 2).

We then treated established *S. aureus* biofilms with sub-MBEC concentrations of fluoro-TEMPO or fluoro-TEMPOMe, stained the treated biofilms for live/dead cells, and examined them using confocal laser scanning microscopy (CLSM). The spectral properties of fluoro-TEMPO and fluoro-TEMPOMe were found to be compatible with the commercially available stains SYTO9 and propidium iodide (PI) (LIVE/DEAD™ bacterial viability kit), with minimal to no interference from spectral overlap from the multi-dye system. Fluoro-TEMPO was found to penetrate the EPS of *S. aureus* ATCC 29213 biofilms, entering cells residing both at the biofilm surface and at the base. Upon cell entry, the fluorescence of fluoro-TEMPO became activated and its presence was clearly visible (Figure 3). Intriguingly, fluoro-TEMPOMe did appear to enter biofilm-residing cells, instead, it appeared to be localized within the EPS of the biofilm (Figure 4). These results suggest that the nitroxide free radical may facilitate EPS penetration and cell entry.

**Figure 3:**
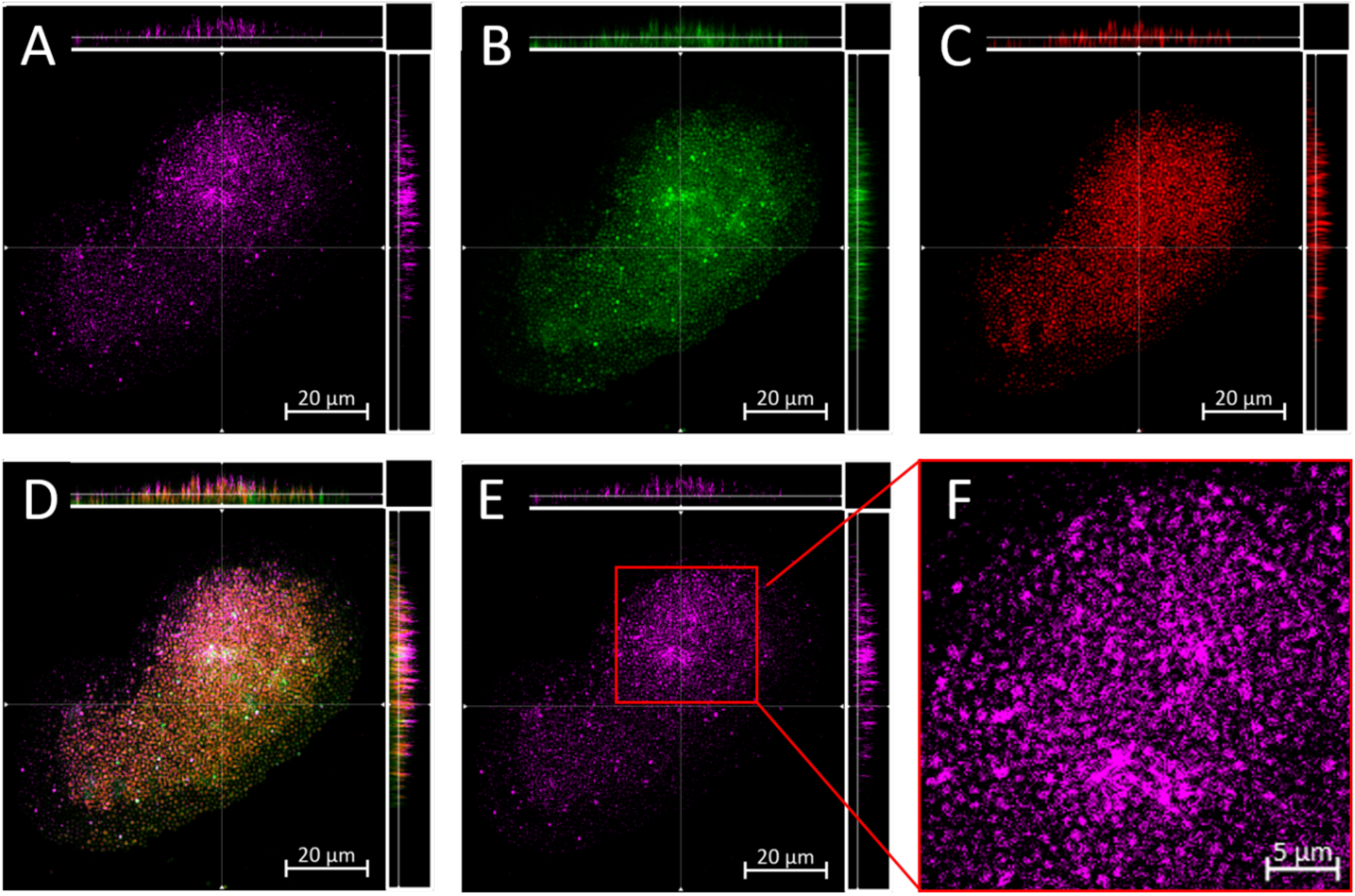
CLSM images of established *S. aureus* ATCC 29213 biofilms treated with fluoro-TEMPO (150 µM, sub-lethal concentration) for 2 hours, then stained with SYTO9, and PI (see methods). (A, E) Excited with 720 nm multi-photon laser (fluoro-TEMPO, pink); (B) Excited with 488 nm laser (SYTO9, live cells, green); (C) Excited with 561 nm laser (PI, dead cells, red); (D) Overlay of images A-C, light grey cells result from merging of green and pink; (F) Expanded image of panel E. The scale bars are 20 µm in length for panels A-E, and 5 µm for panel F. Micrographs show representative horizontal (xy) sections collected within each biofilm, with A-E also showing vertical sections representing the yz and xz planes, shown to the right and top of each individual panel, respectively, taken at the positions indicated by the lines.

**Figure 4:**
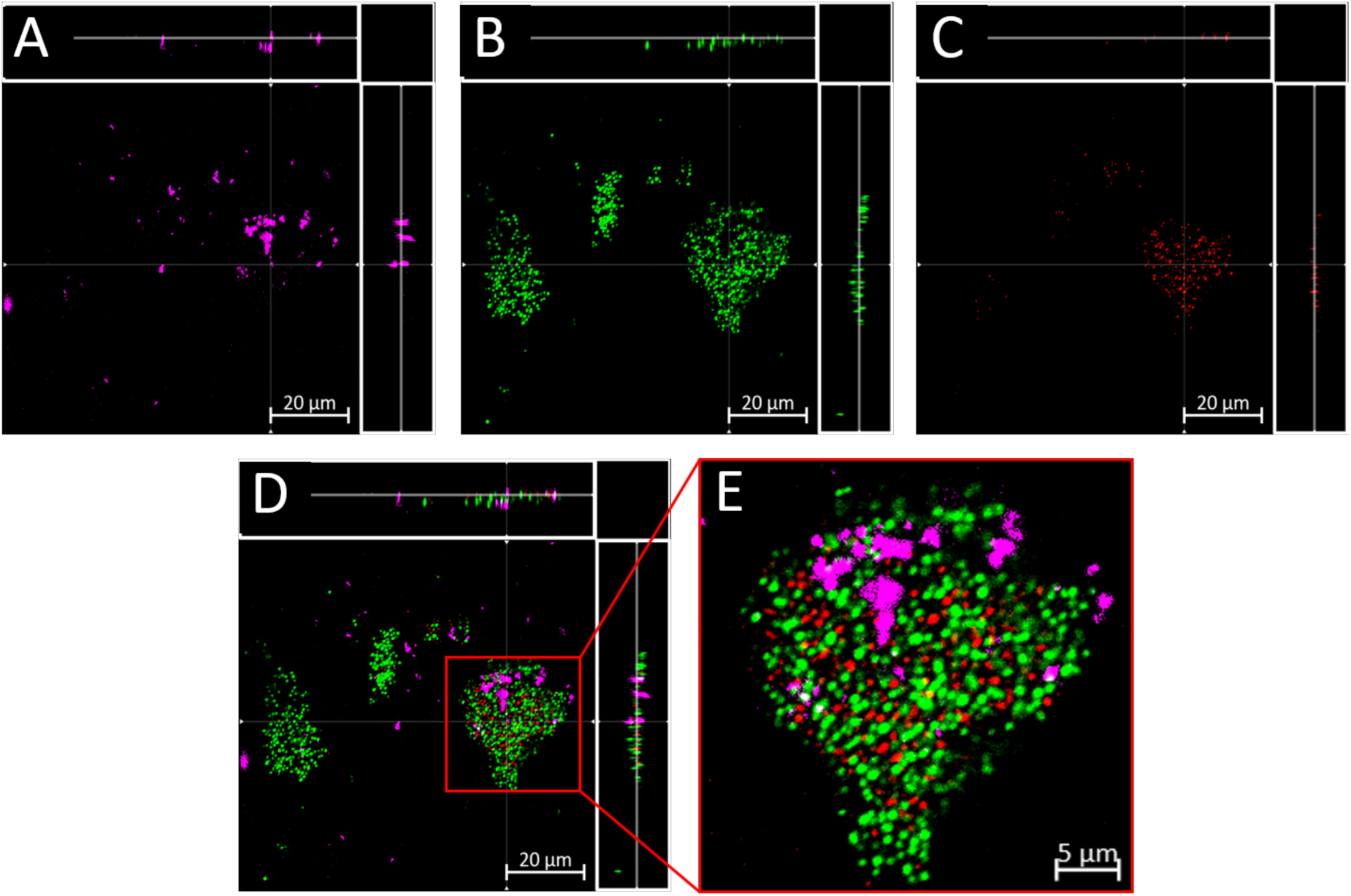
CLSM images of established *S. aureus* ATCC 29213 treated with fluoro-TEMPOMe (150 µM) for 2 hours, then stained with SYTO9, and PI (see methods). (A) Excited with 720 nm multi-photon laser (Fluoro-TEMPOMe, pink); (B) Excited with 488 nm laser (SYTO9, live, green); (C) Excited with 561 nm laser (PI, dead, red); (D) Overlay of images A-C; (E) Expanded image of D. The scale bars are 20 µm in length for images A-D, and 5 µm for image E. Images A-D also show the xy, yz, and xz dimensions.

### Fluoro-TEMPO and CTEMPO are non-toxic to human cells

Our findings suggest that antibiotic-nitroxide hybrids such as fluoro-TEMPO have potential in future clinical applications. To further demonstrate their suitability as preclinical candidates we examined their impact on mammalian cell viability. Fluoro-TEMPO and CTEMPO were evaluated for cytotoxicity against human epithelial T24 cells following a 24-hour cell exposure using the lactate dehydrogenase (LDH) assay. Both fluoro-TEMPO and CTEMPO were found to be non-toxic to T24 cells at concentrations ranging from 20 µM to 720 µM (IC_50_ >720 µM), suggesting they are promising candidates for clinical development.

## DISCUSSION

Currently, treatment options for *S. aureus* biofilm-related infections are limited, with conventional antibiotics often exhibiting little to no therapeutic effect against biofilm-residing bacteria (5, 6). Consequently, treatment for biofilm infections usually involves prolonged high doses of antibiotics and/or surgical removal of infected tissue or implanted medical devices (28). Accordingly, methods for improving the biofilm-eradication activity of conventional antibiotics are urgently needed. Recent studies have demonstrated that nitroxides can mediate biofilm dispersal which can increase the efficacy of the commonly prescribed antibiotic ciprofloxacin against bacterial biofilms (23-25). However, until now, the anti-biofilm properties of nitroxides have only been demonstrated against two Gram-negative pathogens *P. aeruginosa* and *E. coli* (21-25). In this study, we showed that biofilms of the clinically important Gram-positive pathogen *S. aureus* are also susceptible to the biofilm-dispersing properties of nitroxides.

Our findings demonstrate that nitroxide-mediated biofilm dispersal of the Gram-positive pathogen *S. aureus* occurs over a µM concentration range (5 - 80 µM), which is similar to our previous findings for Gram-negative pathogens *P. aeruginosa* and *E. coli* (20 µM) (21, 23). While the precise biofilm dispersal mechanism by nitroxides is currently unknown, nitroxides are considered NO mimics and as such their biofilm-dispersal action potentially involves inhibition of intracellular levels of cyclic di-GMP, which has been directly linked to NO-mediated biofilm dispersal in several Gram-negative and Gram-positive bacterial species (14, 29, 30). The signalling nucleotide cyclic di-GMP has been shown to play a pivotal role in biofilm formation and control, with higher levels facilitating biofilm formation and lower levels triggering biofilm dispersal (31-33). Interestingly, our finding that the optimal nitroxide concentration for biofilm dispersal is similar between Gram-positive and Gram-negative species may suggest that the mechanism by which nitroxide-mediated dispersal occurs is similar between these species.

Our study demonstrates that the nitroxide CTEMPO potentiates the activity of ciprofloxacin against *S. aureus* biofilms, significantly improving its eradication efficacy (>16-fold MBEC improvement). This is consistent with our previous results using nitroxide/ciprofloxacin combination treatment of established *P. aeruginosa* and *E. coli* biofilms (23). Interestingly, other studies which have utilized NO (instead of nitroxides) in combination with antimicrobials have produced similar improvements in antimicrobial activity (e.g. 20-fold increase in chlorine’s activity against multi-species biofilms) (13, 17) further supporting a similarity between the mechanism of NO- and nitroxide-mediated biofilm dispersal. Importantly, the ability of nitroxides to enhance the biofilm-eradication activity of antimicrobials against biofilms is also comparable to other leading methods such as the use of D-amino acids (8-fold improvement over antibiotic alone) (34) and *Cis*-2-decenoic acid (>4-fold improvement over antibiotic alone) (35). Taken together these results support the use of nitroxides as effective enhancers of antibiotic biofilm-eradication activity.

As an improvement to co-treatment, a currently emerging alternative has been the development of dual-acting hybrid compounds. Such compounds combine the anti-biofilm activity of a dispersal agent with an antimicrobial agent to produce a hybrid compound which can both disperse and eradicate biofilm-residing cells. We have previously utilized this strategy to produce ciprofloxacin-nitroxide hybrids which exhibited potent *P. aeruginosa* and *E. coli* biofilm-eradication activity (24, 25, 36). In this study, we have shown that some of these ciprofloxacin-nitroxide hybrids are also potent *S. aureus* biofilm-eradication agents. Importantly, cipro-TMIO was able to completely eradicate (99.9%) established *S. aureus* biofilms at a concentration of only 64 µM (MBEC ≤ 64 µM), which is very similar to its *P. aeruginosa* biofilm-eradication activity (94% eradication at 20 µM) (25). Furthermore, cipro-TMIO’s biofilm-eradication activity is comparable to other promising compounds currently developed as *S. aureus* biofilm-eradication agents, such as halogenated phenazines (MBEC ≤ 10 µM) (37) and quaternary ammonium compounds (MBEC ≤ 25 µM) (38).

Interestingly, in the case of cipro-TMIO the addition of the nitroxide moiety to ciprofloxacin’s core structure does not appear to negatively impact the hybrid’s activity against planktonic *S. aureus* cells (MIC ≤ 1.5 µM, same as ciprofloxacin). However, this trend was not evident in our previous study were cipro-TMIO was administered to *P. aeruginosa* planktonic cells (MIC ≤ 160 µM, > 100-fold increase over ciprofloxacin) (25). These results suggest that the presence of the nitroxide at the secondary amine of ciprofloxacin does not interfere with the fluoroquinolone’s mode of action (inhibition of DNA gyrase and/or topoisomerase IV) in *S. aureus* cells. Thus, it appears that ciprofloxacin-nitroxide hybrids produced via functionalization at the secondary amine of ciprofloxacin may be more suitable for the treatment of *S. aureus* as opposed to *P. aeruginosa* infections. Also, cipro-TMIO’s activity against both planktonic and biofilm-residing *S. aureus* cells as well as its lack of toxicity to human cells (25) makes it an ideal candidate for the treatment of a large number of *S. aureus*-related infections.

The profluorescent nitroxide fluoro-TEMPO is a biofilm-eradication agent with the same anti-biofilm mechanism as cipro-TMIO. Thus, by utilizing the probe properties of fluoro-TEMPO, we demonstrated that the state of the nitroxide (free radical) remains unchanged while interacting with the EPS of the biofilm (i.e., the fluorescence of fluoro-TEMPO is not ‘switched on’ while in the EPS or prior to cell entry). However, as fluoro-TEMPO enters the intracellular space, its fluorescence is quickly activated, indicating that the free radical nitroxide has undergone a chemical change. Furthermore, as our results indicate that the presence of the free radical nitroxide is fundamental to the biofilm-eradication activity of fluoro-TEMPO, it can be inferred that (a) the free radical must be involved in the anti-biofilm activity of nitroxides and (b) the anti-biofilm role of nitroxides must occur via interference and/or regulation of an intracellular process. These findings are in support of the hypothesis that nitroxides may mimic NO and regulate the intracellular levels of c-di-GMP.

Hence, the results presented here support the hypothesis that much like co-treatment (nitroxide/antibiotic), the biofilm-eradication activity of nitroxide-antibiotics, such as cipro-TMIO and fluoro-TEMPO, also occurs via a dual-action mechanism, were the nitroxide moiety triggers biofilm dispersal, and the antibiotic moiety subsequently eradicates the dispersed cells. This dispersal and eradication cycle may initially occur at the surface of the biofilm ultimately exposing previously protected biofilm-residing cells to the antibiotic and allow the process to be repeated until the biofilm is completely eradicated (Figure 5).

**Figure 5:**
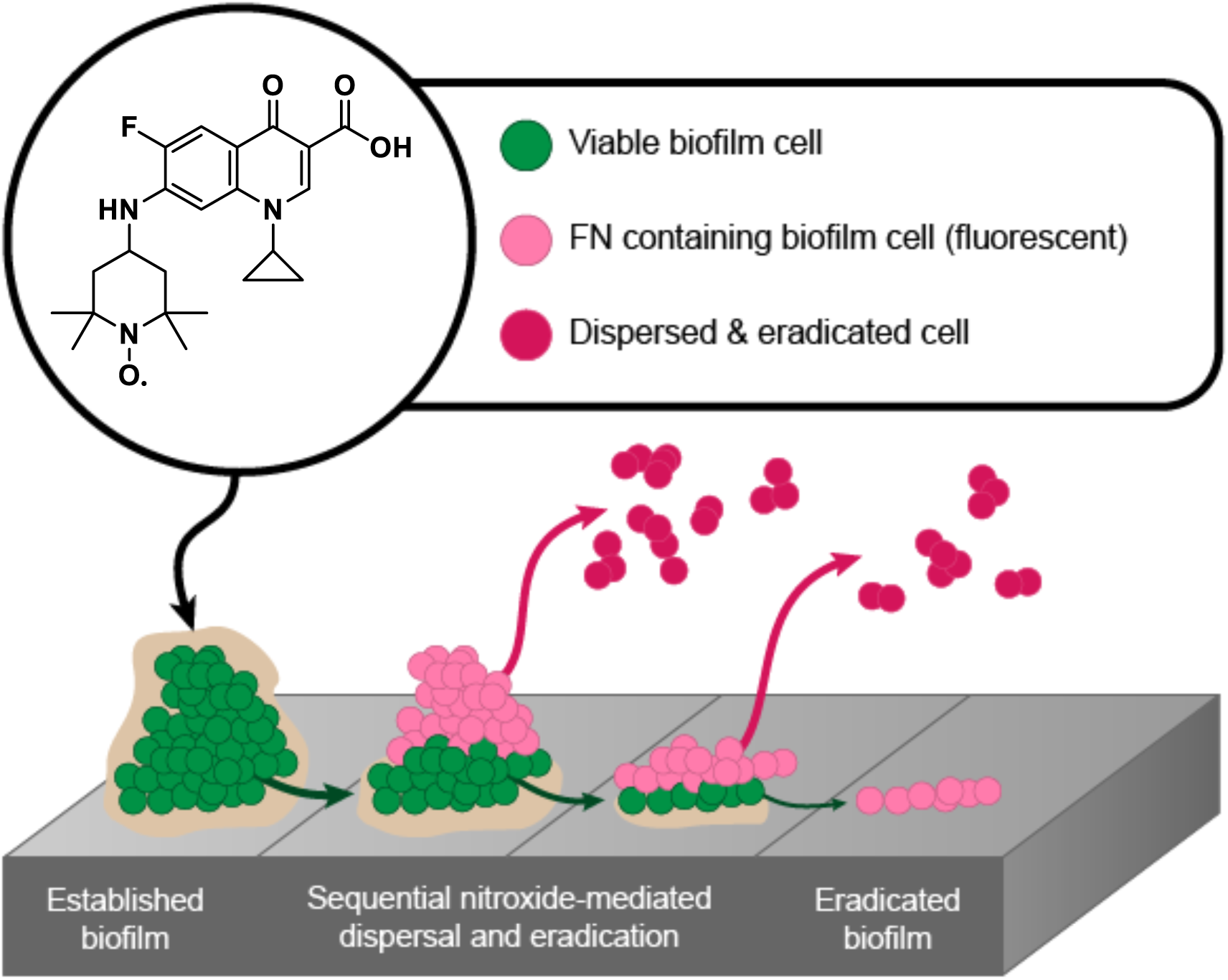
Proposed mechanism of dual-action of fluoro-TEMPO hybrid compound against *S. aureus* biofilms. Upon administration, fluoro-TEMPO enters surface-residing biofilm cells and induces their dispersion from the biofilm. Dispersed fluoro-TEMPO containing cells are subsequently eradicated, which exposes the next layer of biofilm-residing cells. Fluoro-TEMPO enters the newly exposed cells and the process is repeated until the biofilm is completely eradicated.

In conclusion, we have demonstrated that the biofilm dispersal activity of nitroxides is not limited to Gram-negative pathogens but also extends to important Gram-positive pathogens. This work also highlights the ability of nitroxides to restore the antibacterial activity of ciprofloxacin against *S. aureus* biofilms, either by administering them as a combinational treatment or, more potently, as nitroxide functionalized ciprofloxacin derivatives. While our work has focused on the fluoroquinolone class of antibiotics, the methodology followed is easily adaptable to other antimicrobial classes, widely expanding the possible repertoire of anti-biofilm agents that are so urgently needed. Furthermore, our study has provided a promising new therapeutic strategy that could potentially lead to effective broad-spectrum treatments for biofilm-related infections in the not so distant future.

## MATERIALS AND METHODS

### Bacterial strains and culture media

*Staphylococcus aureus* strain ATCC 29213 was used in this study. Bacteria were grown routinely in Lysogeny broth (LB) medium with shaking (200 rpm) at 37 °C. MIC assays were conducted in Mueller Hinton (MH) medium (OXOID, Thermo Fisher, Australia), biofilms were grown in LB medium, and biofilm challenges (dispersal and antimicrobial susceptibility testing) were performed in MH medium or M9 minimal medium (pH 7.0) containing 90 mM Na_2_HPO_4_, 22 mM KH_2_PO_4_, 9 mM NaCl, 19 mM NH_4_Cl, 2 mM MgSO_4_, 100 µM CaCl_2_, and glucose at 22 mM.

### Mammalian cell culture

T24 (ATCC^®^ HTB-4 ™) human bladder epithelial cells, obtained from ATCC were cultured in McCoy’s 5A modified medium (Thermo, Australia) supplemented with 10% heat-inactivated fetal bovine serum (Thermo, Australia). Confluent monolayers were formed by cell culture at 37 °C in a humidified atmosphere of 5% CO_2_.

### Nitroxide and antibiotic stock solutions

The nitroxide 4-carboxy-2,2,6,6-tetramethylpiperidine-1-oxyl (CTEMPO) (Sigma, Australia) was prepared in DMSO at a concentration of 325 mM (stock solution). The antibiotic ciprofloxacin (Sigma, Australia) was prepared in aqueous hydrochloric acid (0.1 M) at a concentration of 60.35 mM (stock solution). Cipro-PROXYL, cipro-TEMPO, cipro-TMIO, fluoro-TEMPO, and fluoro-TEMPOMe were synthesized in-house utilizing previously established procedures (25, 27), and stock solutions were prepared in DMSO (8 mM). All stock solutions were stored in the absence of light at −20 °C. Working solutions were prepared in either M9 minimal media or MH media and used the same day.

### Nitroxide and antibiotic MIC assays

The MIC for CTEMPO, ciprofloxacin, cipro-PROXYL, cipro-TEMPO, cipro-TMIO, fluoro-TEMPO, and fluoro-TEMPOMe were determined by the broth microdilution method, in accordance with the 2015 (M07-A10) Clinical and Laboratory Standards Institute (CLSI). Specifically, in a 96-well plate, twelve two-fold serial dilutions of each compound were prepared to a final volume of 100 µL in MH medium. At the initial time of inoculation, each well was inoculated with 5 × 10^5^ CFU, which had been prepared from fresh overnight cultures. MIC was defined as the lowest concentration of a compound that prevented visible bacterial growth after 18 hours of static incubation at 37 °C (lack of visible growth was further confirmed by spectrophotometric analysis at 600 nm). CTEMPO was tested in the concentration range of 3200 to 1.56 µM, ciprofloxacin in the concentration range of 160 to 0.048 µM, and cipro-PROXYL, cipro-TEMPO, cipro-TMIO, fluoro-TEMPO, fluoro-TEMPOMe were all tested in the concentration range of 1200 to 0.3 µM. Negative controls containing DMSO (vehicle control for CTEMPO, cipro-PROXYL, cipro-TEMPO, cipro-TMIO, fluoro-TEMPO, and fluoro-TEMPOMe) were prepared and serially diluted as above. MIC values for CTEMPO, ciprofloxacin, cipro-PROXYL, cipro-TEMPO, cipro-TMIO, fluoro-TEMPO, and fluoro-TEMPOMe were obtained from at least 2 independent experiments, each consisting of 3 biological replicates and a minimum of 2 technical replicates.

### Biofilm culture in the Calgary Biofilm Device (CBD)

Biofilms were grown and established using the CBD (MBEC^TM^, Innovotech Inc., Canada) used unmodified. The device consists of a two-part reaction vessel. The top component contains 96 identical pegs protruding down from the lid, which fits into a standard flat bottom 96-well plate (bottom component). Biofilm culture was performed as previously described (39). Briefly, overnight LB bacterial cultures were diluted to 10^6^ CFU mL^−1^ in LB and used to inoculate the enclosed flat bottom 96-well plate with ∼10^5^ bacterial cells (130 µL) in each well. The peg lid was inserted into the inoculated wells, and the complete CBD was incubated in a shaking incubator at 150 rpm, 37 °C, and 95% relative humidity for 24 hours.

### Co-treatment (ciprofloxacin/CTEMPO) and antibiotic MBEC assays

Biofilms were established as detailed above. For MBEC value determination, the CBD lid containing established biofilms was removed and rinsed for 10 seconds in PBS (96-well plate, 200 µL in each well), to remove loosely adherent bacteria. The rinsed CBD lid was then transferred to a new flat bottom 96-well plate (challenge plate), which contained the specific treatment. For co-treatment (ciprofloxacin/CTEMPO) experiments the challenge plate contained 2-fold serial dilutions of ciprofloxacin (concentration range 4096 - 32 µM) and CTEMPO at a consistent concentration in each well of either 80, 40, 20, 10, 8, or 2.5 µM in M9 minimal medium (total volume 200 µL per well). For antibiotic MBEC determination, the challenge plate contained 2-fold serial dilutions with a concentration range of 4096 - 32 µM for ciprofloxacin, and a concentration range of 1024 - 8 µM for cipro-PROXYL, cipro-TEMPO, cipro-TMIO, fluoro-TEMPO, or fluoro-TEMPOMe. The complete CBD was then incubated at 37 °C, 120 rpm, 95% relative humidity for 24 hours. The lid was removed from the challenge plate and rinsed twice for 30 seconds in PBS (96-well plate, 200 µL in each well). The rinsed CBD lid, with attached pegs and treated biofilms, was transferred to a new 96-well plate containing fresh LB recovery media. To assist the transfer of any remaining viable cells to the recovery media, the device was sonicated for 30 minutes (<20 °C). The peg lid was then discarded, and the biofilm-recovered bacteria from each well of the recovery plate was serially diluted in PBS, and then triplicate 5 µL aliquots of each dilution were plated onto LB agar and incubated at 37 °C overnight for viable CFU enumeration. MBEC values were determined as the lowest concentration that resulted in either zero growth or a CFU value ≤ 99.9% of the untreated controls. MBEC values were obtained from at least 2 independent experiments, each consisting of 3 biological replicates and 3 technical replicates minimum.

### Nitroxide-mediated biofilm dispersal assays

Biofilm dispersal assays were performed as described above for MBEC assays, with the following modification and additional steps. The challenge plate contained 2-fold serial dilutions of CTEMPO (concentration range 160 - 2.5 µM) and ciprofloxacin at a consistent concentration of 6 µM in M9 minimal medium (total volume 200 µL per well), control wells contained ciprofloxacin only at 6 µM. After sonication, the peg lid was discarded, and the recovery plate containing biofilm-recovered bacteria was covered with a breathable sealing membrane (Breathe-Easy^®^ sealing membrane, Sigma, Australia), and incubated at 37 °C for 20 hours with shaking in microtiter plate reader (BMG, Australia). OD_600_ measurements were obtained every 15 minutes over the 20 hour period. Growth curves for each well, containing the recovered biofilm cells, were then generated and a non-linear regression function was applied to determine the lag time of each sample (lag time is directly proportional to the initial CFU count of each well, with wells initially containing a low CFU counts producing a longer lag time than wells with a higher initial CFU count) (40, 41).

### LDH release cytotoxicity assay

The cytotoxicity of fluoro-TEMPO and CTEMPO against human T24 urinary bladder epithelial cells was examined utilizing the Pierce™ LDH cytotoxicity assay kit (Life Technologies, Australia) as per manufacturer’s instructions. Briefly, triplicate confluent T24 cell monolayers were treated with fluoro-TEMPO or CTEMPO (concentrations between 720 - 20 µM) for 24 hours at 37 °C in a humidified atmosphere of 5% CO_2_. Cells treated with DMSO/PBS (4.5% DMSO final concentration) or sterile water served as negative controls and cells treated with 10X lysis buffer (maximum LDH release) served as a positive control. After 24 hours incubation, 50 µL of the supernatant was transferred into a new 96-well plate, mixed with 50 µL of the reaction mixture (LDH assay kit) and incubated at room temperature (protected from light) for 30 minutes before the stop solution (50 µL) was added. The plate was then centrifuged (1000 × *g*) for 5 minutes to remove air bubbles, and the absorbance at 490 and 680 nm was measured in a spectrostar (BMG) plate reader.

### Confocal laser scanning microscopy of *S*. *aureus* biofilms

Biofilms were grown on the CBD as noted above. Established biofilms were treated with fluoro-TEMPO or fluoro-TEMPOMe (150 µM) in M9 medium, incubated at 37 °C for 2 hours, rinsed in PBS for 5 seconds, and stained with LIVE/DEAD™ *Bac*Light™ Bacterial Viability kit L7007 (Life Technologies, Australia) as per manufacture’s protocol. Treated and stained biofilms were mounted using ProLong^®^ Diamond Antifade Mountant (Life Technologies, Australia) and immediately analyzed by CLSM. CLSM was conducted on a Zeiss 780 NLO Point Scanning Confocal, equipped with a Mai-Tai deep see multi-photon laser (tuneable between 690 and 1040 nm). For fluoro-TEMPO and fluoro-TEMPOMe the Mai-Tai laser was set at 720 nm (a wavelength which did not excite SYTO9 or propidium iodide), for SYTO9 the 488 nm laser was used, and the 561 nm laser was applied for PI. All imaging experiments utilized a 100 × oil immersion objective.

### Calculating lag time for biofilm dispersal experiments

Growth curve time points and corresponding OD_600_ values were exported to an excel spreadsheet, where a non-linear regression was applied using solver. Lag times were determined for each replicate for each condition tested and group medians were compared by a Kruskal-Wallis test (GraphPad Prism 7).

## ACKNOWLEDGMENTS

This work was supported by a Queensland University of Technology (QUT) grant (to M.T and K.E.F.-S.) and an Asian Office of Aerospace Research and Development Grant (FA2386-16-1-4094, R&D 16IOA094). AV is supported by an Australian Government Research Training Program (RTP) Scholarship; RD by a National Health and Medical Research Council grant (APP1144046 to MT); KEF-S by an Australian Research Council Future Fellowship (FT140100746); and MT by a QUT Vice-Chancellor’s Research Fellowship. The authors would also like to thank Professor Flavia Huygens, for the provision of *S. aureus* strains.

## ABBREVIATIONS

CBD: Calgary Biofilm Device
c-di-GMP: bis-(3′-5′)-cyclic dimeric guanosine monophosphate
cipro-PROXYL: 1-Cyclopropyl-6-fluoro-7-(4-(2,2,5,5-tetramethyl-1-oxy-pyrrolidine-3-carbonyl)piperazin-1-yl)-4-oxo-1,4-dihydroquinoline-3-carboxylic acid
cipro-TEMPO: 1-Cyclopropyl-6-fluoro-7-(4-(2,2,6,6-tetramethyl-1-oxy-piperidine-4-carbonyl)piperazin-1-yl)-4-oxo-1,4-dihydroquinoline-3-carboxylic acid
cipro-TMIO: 1-Cyclopropyl-6-fluoro-7-(4-(1,1,3,3-tetramethylisoindolin-2-yloxyl-5-carbonyl)piperazin-1-yl)-4-oxo-1,4-dihydroquinoline-3-carboxylic acid
CFU: colony forming units
CTEMPO: 4-carboxy-2,2,6,6-tetramethylpiperidin-1-yloxyl
DMSO: dimethyl sulfoxide
EPS: extracellular polymeric substances
fluoro-TEMPO: 1-Cyclopropyl-6-fluoro-7-(2,2,6,6-tetramethyl-1-oxy-piperidine-4-yl)amino)-4-oxo-1,4-dihydroquinoline-3-carboxylic acid
fluoro-TEMPOMe: 1-Cyclopropyl-6-fluoro-7-((1-methoxy-2,2,6,6-tetramethylpiperidine-4-yl)amino)-4-oxo-1,4-dihydroquinoline-3-carboxylic acid
MBC: minimal bactericidal concentration
MBEC: minimal biofilm eradication concentration
MIC: minimal inhibitory concentration
NO: nitric oxide

